# BEREN: A bioinformatic tool for recovering Giant viruses, Polinton-like Viruses, and Virophages in metagenomic data

**DOI:** 10.1101/2024.10.09.617401

**Authors:** Benjamin Minch, Mohammad Moniruzzaman

## Abstract

Viruses in the kingdom *Bamfordvirae*, specifically giant viruses (NCLDVs) in the phylum *Nucleocytoviricota* and smaller members in the *Preplasmiviricota* phylum, are widespread and important groups of viruses that infect eukaryotes. While viruses in this kingdom such as giant viruses, polinton-like viruses, and virophages have gained large interest from researchers in recent years, there is still a lack of streamlined tools for the recovery of their genomes from metagenomic datasets. Here, we present BEREN, a comprehensive bioinformatic tool to unlock the diversity of these viruses in metagenomes through five modules for NCLDV genome, contig, and marker gene recovery, metabolic protein annotation, and *Preplasmiviricota* genome identification and annotation. BEREN’s performance was benchmarked against other mainstream virus recovery tools using a mock metagenome, demonstrating superior recovery rates of NCLDV contigs and *Preplasmiviricota* genomes. Applied to a real-world dataset from the Baltic Sea, BEREN identified diverse *Bamfordvirae* members, giving insight into viral interactions and metabolic functions in this region. Overall, BEREN offers a user-friendly, transparent bioinformatic solution for studying the ecological and functional roles of these eukaryotic viruses, facilitating broader access to their metagenomic analysis.

## Introduction

Eukaryotic viruses are widespread and important members of the global virosphere, infecting both charismatic macrofauna such as pigs and smaller microeukaryotes like photosynthetic algae. One of the major kingdoms of these viruses is *Bamfordvirae*, which includes the *Nucleocytoviricota* phylum (giant viruses) [1] and the closely related *Preplasmiviricota* phylum which contains virophages, polintons, and polinton-like viruses [2]. Recent research has highlighted the widespread presence of viruses from these two phyla of double-stranded DNA viruses in diverse ecosystems, including both terrestrial and aquatic environments [3], [4], [5], [6]. These viruses also often infect environmentally important microeukaryotic hosts such as the bloom-forming algae *Emiliania huxleyi*, and *Phaeocystis globosa* [7], [8], [9].

Giant viruses have large genomes (up to 2.5Mbp) and particle size (up to 2uM)[10], [11], and the ability to influence host metabolism through a suite of metabolic genes [12], [13], [14], [15]. In addition to this, recent studies have also revealed that virophages within *Preplasmiviricota* are key antagonists of giant viruses within the *Nucleocytoviricota*, and could be adopted as defense mechanisms against giant viruses by certain protists as integrated elements in their genomes [16]. These viruses are typically much smaller in size (<0.2uM) and have much smaller genome sizes ranging typically from 10-50kbp [17]. Given their environmental distribution and intriguing relationships, there has been increasing interest regarding their diversity, ecological dynamics and host associations [16], [17], [18].

While many bioinformatic tools exist for recovering viruses from metagenomes, most of these tools have a strong emphasis on identification of prokaryotic viruses such as phages infecting bacteria [19]. Some recent tools such as geNomad and IPEV [20], [21] can identify dsDNA eukaryotic viruses, but these tools often lack specificity as to the identity of the eukaryotic virus, or the ability to recover full genomes of these viruses for comprehensive downstream analysis. In addition, these tools don’t take into account the ever-expanding diversity of these viruses such as the newly discovered *Mirusviricota* phylum [22], Egovirales order [23], or Mriyaviricetes class [24].

Currently, a comprehensive tool to identify and analyze members of the *Bamfordvirae* that infect diverse eukaryotes -specifically giant viruses, polinton-like viruses, and virophages does not exist. This presents a major hurdle for researchers looking to gain a holistic understanding of the role of these viruses in their study environment. As a case in point, giant viruses and virophages share deep evolutionary history, and their antagonistic interactions are likely pervasive in diverse environments that potentially have important implications for protist population dynamics [17], [25]. In addition to this, understanding their diversity, metabolic potential and host association necessitates robust identification of these viral genomes. Tools that automate simultaneous identification and characterization of these viruses in diverse datasets will pave the way for new hypotheses regarding the relationship of these viruses and their role in protist population dynamics, and terrestrial and aquatic food webs.

There is currently no consensus methodology for recovery of these viruses [4], [5], [13], [17], [26] and many of the proposed methods require extensive bioinformatic expertise and manual screening. Here we present BEREN, a “one-stop-shop” for uncovering the diversity and metabolic potential of giant viruses, polinton-like viruses, and virophages in any metagenomic sample. This tool both opens up the realm of *Bamfordvirae* viruses to interested researchers and provides a streamlined methodology to increase repeatability. We demonstrate that this tool outperforms available virus recovery tools in recovering NCLDV contigs and *Preplasmiviricota* genomes, as well as provides extra functionality integrated in the pipeline, such as genome binning, taxonomy, decontamination and metabolic annotation. We hope this tool will be used to increase our understanding of these viruses in diverse environments and help democratize the field of eukaryotic virus ecogenomics for scientists without speciality in bioinformatics.

## Methods

### The BEREN tool

The BEREN tool is made up of five different modules that can be run independently or all together on a metagenomic dataset. These modules include (1) NCLDV markers, (2) NCLDV contigs, (3) NCLDV bins, (4) metabolism and protein annotation, and (5) *Preplasmiviricota* identification. The tool is publicly available at: https://gitlab.com/benminch1/BEREN. Detailed instructions and necessary scripts for installation and database downloads are also provided there along with default parameters for the tools integrated in the BEREN workflow.

#### NCLDV marker module

The main purpose of the NCLDV marker module is to gain insight into the full diversity of NCLDVs within a given metagenomic dataset, including those that remain elusive to genome binning methods due to low coverage or abundance, as well as newly discovered NCLDV relatives such as Egoviruses, Mirusviruses, and Mriyaviruses. When run, this module first predicts proteins from the metagenomic assembly using prodigal-gv, a version of prodigal trained with NCLDV genomes [20], [27]. It then searches for NCLDV marker genes with the NCLDV_markersearch script [1]. The marker genes searched for include the NCLDV major capsid protein (MCP), DEAD/SNF2-like helicase (SFII), DNA-directed RNA polymerase beta and alpha subunits (RNAPS and RNAPL), DNA polymerase family B (PolB), Transcription initiation factor IIB (TFIIB), DNA topoisomerase II (TopoII), Packaging ATPase (A32), and the Poxvirus Late Transcription Factor VLTF3 (VLTF3).

After storing all markers in separate protein fasta files, this module will use the newly discovered PolB markers in the target metagenome dataset to build a phylogenetic tree. The sequences are first aligned using MAFFT [28] to a custom set of reference NCLDV PolBs representing all NCLDV families as well as a set of PolBs from bacteria and eukaryotes for quality filtering. The alignment is then trimmed with trimAL using the ‘-gt 0.1’ parameter [29] and a phylogenetic tree is constructed using fasttree [30]. This is all wrapped within the NuPhylo script, available as a separate tool (github.com/BenMinch/Nuphylo).

In addition to providing insight into traditional NCLDVs, the NCLDV marker module can also search for marker genes found in the newly discovered Egovirales order, proposed *Mriyaviricetes* class [24] and *Mirusviricota* phylum. The dataset is searched for *Mirusviricota* and Egovirales MCPs using hmm profiles obtained from Gaïa et al. [23] [22], and *Mryiyaviricetes* MCPs using a custom hmm profile. All of these hmms are leveraged in a custom script for parsing.

#### NCLDV contig module

While NCLDV genome recovery is an obvious goal of many researchers wanting to study these viruses, recovery of full genomes might not be always possible due to fragmented assemblies and known chimerism of NCLDVs [31]. For these reasons, we have included an NCLDV contig module, which allows users to look at NCLDVs from both a genomic and contig perspective. NCLDV contigs are often used similarly to microbial operational taxonomic units (OTUs) in that they can be used to create abundance profiles and gain functional insight on the NCLDVs present in a metagenomic sample [32]. In the NCLDV contig pipeline, NCLDV contigs are retrieved using ViralRecall [33], which identifies NCLDV contigs based on homology to known NCLDV proteins. The identified contigs are screened for an NCLDV score of >=3 and a length of >10kbp. These contigs can be used for downstream profiling of functional potential encoded by the NCLDVs in the target dataset.

#### NCLDV bins module

This module is designed to recover both high-quality and partial NCLDV genomes through a genome binning, screening, and cleaning pipeline. First, metagenomic contigs are binned using metabat2 [34], and then potential NCLDV bins are identified using the NCLDV markersearch script [1] after proteins are predicted using prodigal-gv[20]. Bins with >=1 hit to an NCLDV marker gene are retained for further analysis. These bins are then screened using ViralRecall in ‘contig’ mode [33] to confirm they are NCLDVs. Bins with negative ViralRecall scores are further investigated to assess whether they represent partial NCLDV genomes.

After this screening, another screening is performed to further separate “high-quality” and “partial” NCLDV bins. High-quality NCLDV bins are defined as having at least 3 of the following marker genes (PolB, MCP, A32, VLTF3, SFII) so they can be easily used in a concatenated phylogeny [1]. Both partial and high-quality bins are then processed using ViralRecall in ‘contig’ mode, which screens individual contigs within the bins to remove potential cellular contamination. Specifically, contigs that have a negative ViralRecall score are discarded from the bins. Taxonomy for these bins is assigned using TIGTOG, a tool that predicts taxonomy through machine learning approach [35].

This module also has the possibility of recovering some Mirusviricota and majority of the Mriyavirus genomes in a metagenomic dataset as these viruses share many of the same marker genes and orthologous groups with their NCLDV counterparts [22], [24]. Through testing, these bins typically end up being classified as partial as they have lower ViralRecall scores compared to NCLDVs and fewer identified marker genes (Figure S1a). A simple test using a mock metagenome assembly containing 111 Mirusvirus and 60 Mriyavirus genomes demonstrated BEREN was able to recover 31% of Mirusvirus genomes and 91% of Mriyavirus genomes (Figure S1b). These bins are flagged by BEREN and put into separate folders for further analysis at the discretion of the researcher.

#### Metabolism and protein annotation module

Many NCLDVs encode genes for a wide range of cellular metabolic processes, potentially enabling them to reprogram cellular metabolism in myriad different ways, which is unprecedented in the virus world. The metabolism and protein annotation module attempts to uncover these unique functions of NCLDVs through hmm-based protein annotation using multiple databases. Using recovered NCLDV contigs, NCLDV high-quality bins, and partial bins, this module first annotates proteins using Pfam [36], GVOG [1], and KEGG [37] databases. Metabolic proteins of interest are parsed into a separate file using a custom script (github.com/BenMinch/nump). Resulting protein annotations can be used to analyze the functional profiles of the whole NCLDV community or individual NCLDV genomes.

#### Preplasmiviricota module

Virophages, Polintons, and Polinton-like viruses (PLVs) are all part of *Preplasmiviricota*, a sister phyla to NCLDVs [2]. These smaller eukaryotic viruses are widespread in aquatic ecosystems [5] and often interact with NCLDVs [17], [18], [25], making their recovery important for understanding eukaryotic virus ecology. The *Preplasmiviricota* module effectively recovers and identifies viruses of the *Preplasmiviricota* phylum in metagenomic datasets.

First, contigs potentially belonging to viruses in this phylum are screened using a modified version of ViralRecall with new protein profiles common across diverse *Preplasmiviricota* members [38]. Contigs with a positive score are then screened to only keep contigs with a major capsid protein as well as one other marker gene (integrase, minor capsid protein, and Ftsk ATPase). Taxonomy is assigned to these contigs using the Virophage_affiliation script [17]. Protein annotations for *Preplasmiviricota* contigs are performed using both the Pfam database [36] and a custom hmm profile of environmental *Preplasmiviricota* protein clusters [5].

### Benchmarking BEREN

#### Mock metagenome creation

To test the capabilities of the BEREN tool at recovering diverse NCLDV and *Preplasmiviricota* members, a mock metagenome was created using FetaGenome2 (github.com/OLC-Bioinformatics/FetaGenome2). To create the mock community, 100 genomes of unique members of *Preplasmiviricota* [5] were obtained (dereplicated at 80% ANI using dRep [39]) as well as 69 unique NCLDV genomes from the GOEV database [22](637 total contigs) representing 3 from each major NCLDV family, covering all NCLDV orders (Imitervirales, Algavirales, Pimascovirales, Pandoravirales, Chitovirales, Asfuvirales). Ten bacterial genomes were also used to simulate ubiquitous bacterial populations within environmental metagenomic samples, and these genomes were obtained from the GORG tropics database [40]. The mock community was set up so that 80% of the reads would belong to bacterial genomes, 10% to NCLDV genomes, and 10% to *Preplasmiviricota* genomes.

The mock metagenome was created using Illumina paired-end short reads of 250bp mean length. A total of 100 million paired-end reads were simulated. These paired-end reads were then quality-trimmed using cutadapt [41] and assembled into a mock metagenome using MEGAHIT [42].

#### Benchmarking BEREN and other virus recovery tools

The BEREN tool was benchmarked against a set of other popular virus recovery tools including DeepVirFinder [43], geNomad [20], DeepMicroClass [44], VIBRANT [45], ViralVerify [46], Virsorter2 [47], and IPEV [21]. While most of these tools were built with prokaryotic viruses in mind, they can often perform moderately well at recovering eukaryotic viruses [20].

All tools were run with default settings using the mock metagenome as input. Many of these tools (with the exception of geNomad) will not explicitly discriminate between NCLDV and *Preplasmiviricota* viruses, so sequences identified as viral were run through the BEREN pipeline to parse these two groups apart. For tools using deep learning (DeepVirFinder, DeepMicroClass, and IPEV) a score cutoff of 0.9 was used to determine viral sequences. Percent recovery of NCLDV contigs as well as *Preplasmiviricota* genomes was evaluated for each tool.

#### Benchmarking BEREN for NCLDV genome completeness

To assess the ability of BEREN to recover high-quality “complete” NCLDV genomes, the NCLDV bin module was run on the mock metagenome. Each recovered genome bin was compared to the known genomes in the mock community using BLASTn [48]. Completeness was defined to be the percentage of the total genome length recovered by BEREN.

### Testing BEREN on a real-world metagenome

The BEREN tool was applied to a real-world, publicly available aquatic metagenomic dataset from the Baltic Sea [49] to test the ability of the tool to gain insight into the diversity of eukaryotic viruses in this ecosystem. Briefly, 10 sets of paired-end raw read libraries were downloaded from NCBI (Table S1) representing 5 cellular fraction (>0.2uM) and 5 viral fraction (<0.2uM) metagenomes. These reads were trimmed using cutadapt [41] and assembled using MEGAHIT [42]. The NCLDV markers, contigs, bin, and metabolism modules were run on the cellular fraction metagenome and the *Preplasmiviricota* module was run on the viral fraction metagenome. All analysis and graphs were done using R and ggplot2 [50] and all phylogenetic trees were visualized with iTOL [51] or anvi’o [52].

## Results

### BEREN exceeds other automated virus detection tools in NCLDV and *Preplasmiviricota* recovery

A benchmarking of the BEREN tool against other popular viral recovery tools (DeepVirFinder, geNomad, DeepMicroClass, VIBRANT, ViralVerify, Virsorter2, and IPEV) showed BEREN outperforms these tools on both NCLDV contig and *Preplasmiviricota* genome retrieval from the mock metagenome. Out of the 637 NCLDV contigs above 10kbp in the mock metagenome, BEREN was able to recover 583 contigs (92%) (Figure 2a). The next best tool, DeepMicroClass, was able to identify 83.3% of these contigs, with many tools recovering below 50%. Since BEREN’s NCLDV contig recovery relies on ViralRecall, this result is largely attributed to the performance of this tool at identifying NCLDV contigs in metagenomic data.

**Figure 1.**
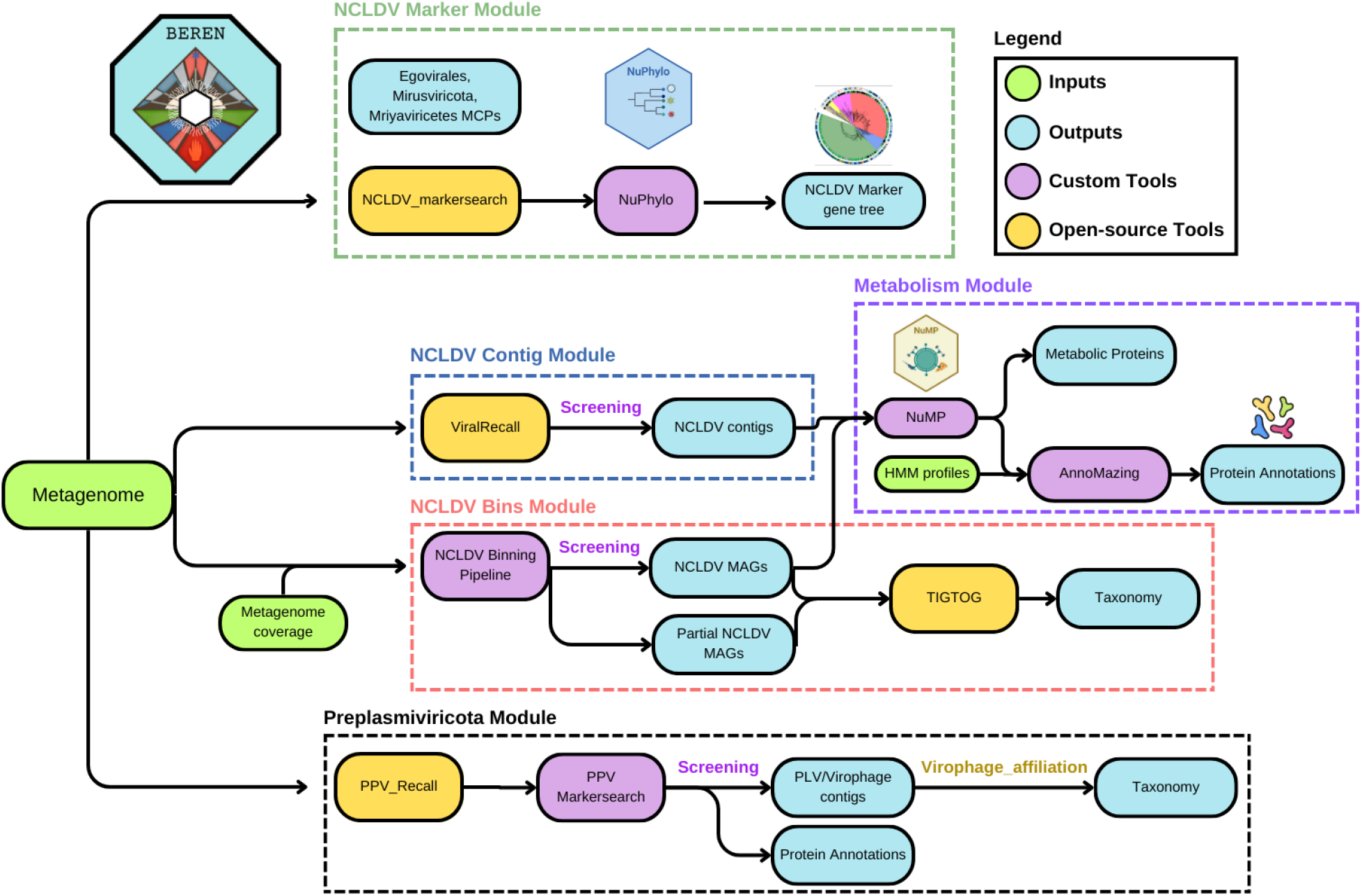
The BEREN pipeline. BEREN consists of 5 major modules, each comprising of multiple open-source tools as well as custom scripts. The input of a metagenome can yield NCLDV markers, contigs, and genomic bins, *Preplasmiviricota* genomes, as well as the metabolic potential and protein annotation of these viruses.

**Figure 2.**
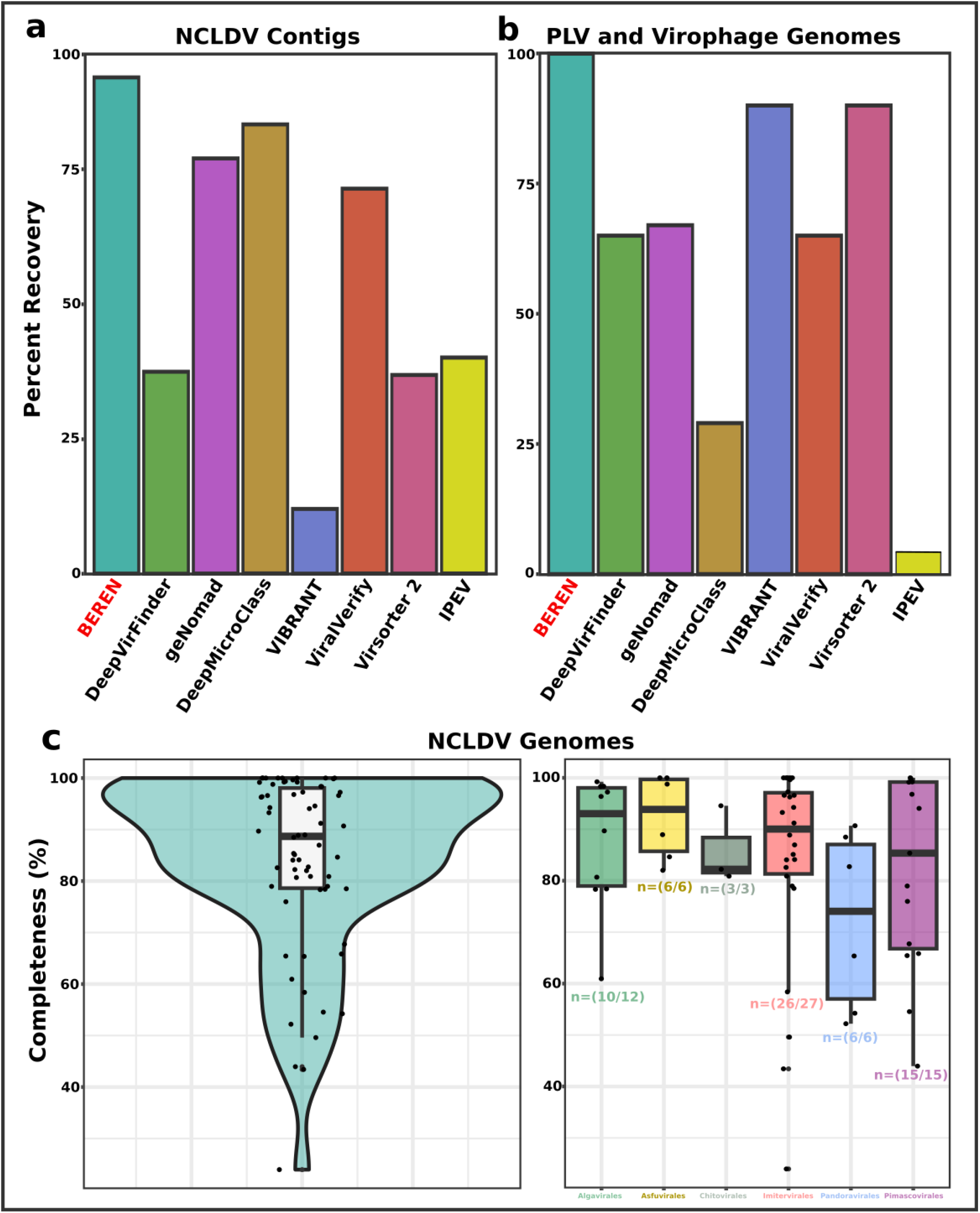
Benchmarking of BEREN against other virus recovery tools. The recovery of **(a)** NCLDV contigs and **(b)** PLV and virophage genomes was benchmarked against popular virus identification tools. Each tool was run with default settings and total recovery is shown here. **(c)** The completeness of the recovered NCLDV genomes from BEREN was tested by comparing recovered genomes to those used to build the mock metagenome. Completeness is defined as the percentage of the total length of the genome recovered. This information is also provided for each NCLDV order with the number of recovered genomes in that order below the boxplots.

Recovery of *Preplasmiviricota* genomes gave similar results as BEREN recovered all (100%) of the 100 genomes seeded into the mock metagenome (Figure 2b). Other tools recovered between 4% and 90% of the *Preplasmiviricota* genomes, with VIBRANT and Virsorter2 performing the best.

### BEREN recovers high-quality NCLDV genomes

Out of the 69 seeded NCLDV genomes, BEREN was able to recover 66 (95.6%). Using BLASTn to match the recovered genomes with the seeded genomes, we were able to calculate the completeness of each recovered genome (percent of genome length recovered). Genomes recovered with BEREN had an average completeness of 84% and a median completeness of 89% (Figure 2c). Genomes recovered within the Pandoravirales order had the lowest average completeness of 72%, while Asfuvirales had the highest average completeness of 92% (Figure 2c).

### BEREN comprehensively assesses the diversity and metabolic potential of eukaryotic viruses in a real-world metagenome

A publicly available metagenomic dataset from the Baltic Sea was downloaded to demonstrate a real-world application of the tool. This dataset consisted of ten metagenomes, five from the cellular fraction (>0.2uM) and five from the viral fraction (<0.2uM). BEREN was run with all NCLDV modules on the cellular fraction and the *Preplasmiviricota* module on the viral fraction.

A total of 482 quality-filtered NCLDV PolB marker genes were recovered from the Baltic datasets (Figure 3a). These represented all major NCLDV orders with the majority being either Algavirales and Imitervirales. From the five cellular fraction samples, 492 NCLDV contigs were recovered ranging from 10-117 kbp (Figure 3b). Many of these contigs contained multiple marker genes with certain contigs containing up to 8. For contigs over 30kbp the mean number of marker genes ranged from 1 to 4. 53 NCLDV bins were also recovered from the 5 cellular metagenomes including 2 Pandoravirales bins, 21 Imitervirales bins, and 30 Algavirales bins (Figure 3c). These genomic bins ranged in size from 52-485 kbp and encoded a variety of metabolic genes such as those involved in light harvesting, DNA processing, and carbon metabolism.

**Figure 3.**
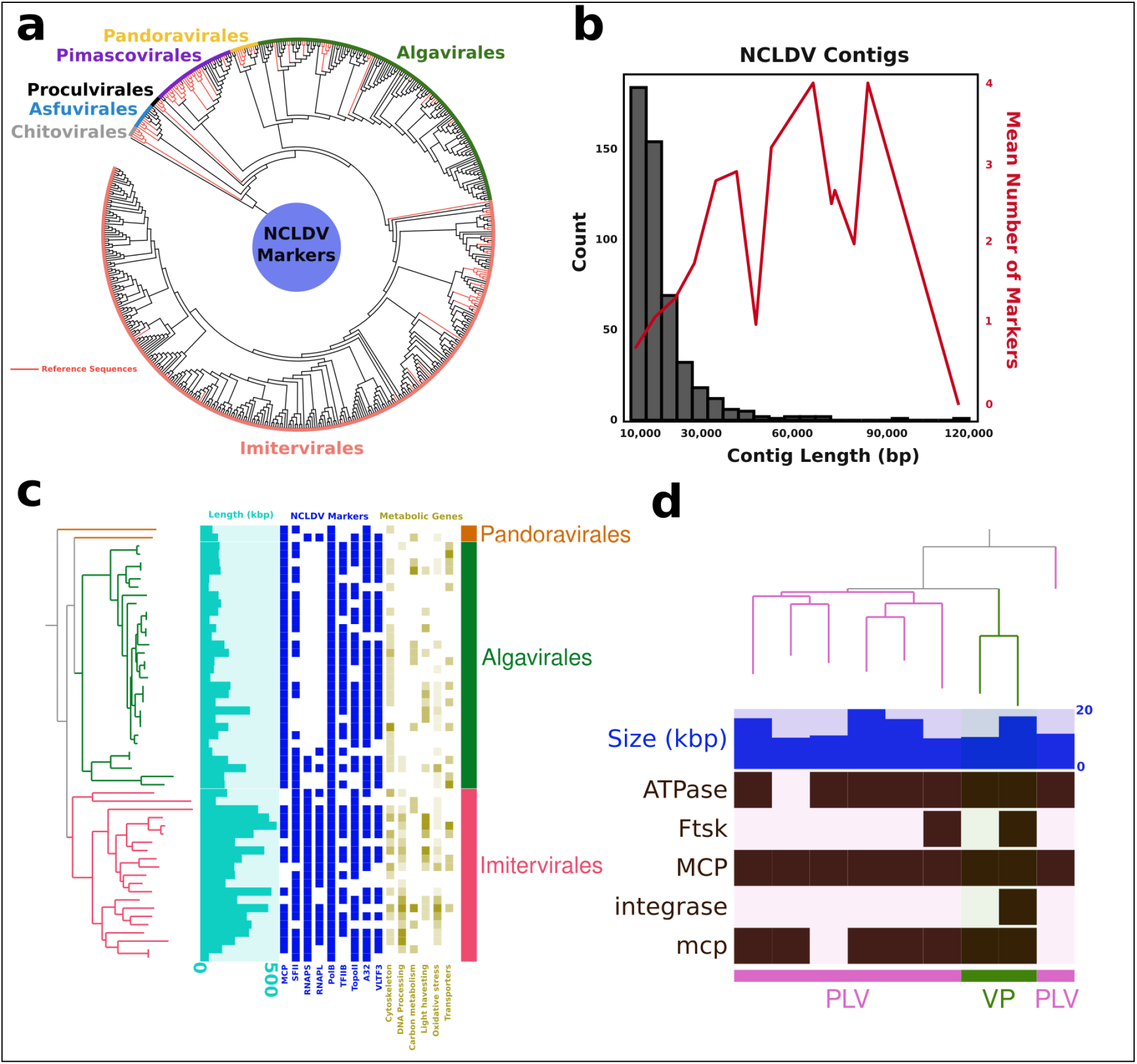
Testing BEREN on real-world Baltic Sea metagenomes. BEREN was used to assess the diversity of NCLDVs and *Preplasmiviricota* in 10 Baltic Sea metagenomes. **(a)** A phylogenetic tree of recovered NCLDV PolB marker genes, clustered with custom reference sequences from the GOEV database (ref). **(b)** NCLDV contigs recovered from the five cellular fraction metagenomes. The red line represents the mean number of NCLDV markers in a given “bin” or size class. **(c)** A phylogeny of recovered NCLDV genomes was created using the PolB marker gene. Genome length, NCLDV markers, and metabolic gene categories are also represented on this tree. **(d)** *Preplasmiviricota* genomes recovered from the viral fraction metagenomes were phylogenetically placed based on the major capsid protein (MCP). Other marker genes as well as genome size are present in the tree. *Preplasmiviricota* genomes are classified as Polinton-like virus (PLV) or Virophage (VP) by BEREN.

After running BEREN on the five viral-fraction samples, a total of 9 viruses of the phylum *Preplasmiviricota* were recovered (Figure 3d). These viruses ranged in size from 10.5-19.9 kbp and all encoded a major capsid protein as well as another marker gene. Using BEREN, two of these genomes were identified as Virophages, while the rest were most likely Polinton-like viruses (PLVs).

## Discussion

In an age with petabytes of sequencing data publicly available on NCBI, streamlined pipelines are needed to enable robust discovery of both established and emerging, novel diversity of viruses in high-throughput sequencing datasets . For eukaryotic viruses belonging to the *Nucleocytoviricota* and *Preplasmiviricota* phyla, BEREN provides an easy-to-use framework for researchers to start looking into the diversity and metabolic potential of these viruses in all environments.

In addition to outperforming other viral identification tools in the realm of NCLDVs and *Preplsmiviricota* recovery, BEREN has several other advantages. One of such advantages is that BEREN is designed specifically for these two groups of viruses, and therefore provides many helpful downstream outputs for studying their diversity - such as marker genes for phylogenetic and taxonomic assignment. While several other tools were able to recover NCLDVs and *Preplasmiviricota* viruses, many of them do not provide any distinguishing taxonomic or phylogenetic information about these viruses. For example, DeepMicroClass and IPEV label both of these phyla as “Eukaryote Virus” and other tools like VIBRANT, DeepVirFinder, and Virsorter2 don’t distinguish them from bacterial viruses. This can be partially attributed to the recent and ongoing research surrounding *Preplasmiviricota* phylogeny and taxonomy [2], which we sought to leverage in our tool.

Another advantage of BEREN is that it is able to recover NCLDV genomic bins in addition to contigs. As far as we know, no other virus-identification tool tested can recover NCLDV genomic bins, which is a key limitation as NCLDV genomes are often too large to be represented by a single contig [1]. Getting genomic bins also has the advantage of providing genome annotations to uncover the functional potential of these viruses. Other virus tools we tested either do not give this information, or use databases tailored to phages [45].

A third advantage of BEREN is that it does not rely on neural network methods, where intermediate steps in the pipeline are often not easily interpretable. While neural network-based methods like DeepMicroClass and DeepVirFinder offer powerful capabilities, BEREN provides additional flexibility by allowing users to adjust parameters to relax or constrain virus searches. BEREN’s methodology is transparent and easily interpretable by researchers. This transparency allows for easy interpretation and modification of BEREN to suit specific research needs.

## Conclusion

Overall, BEREN represents a “one-stop shop” for researchers interested in studying NCLDV and *Preplasmiviricota* viruses in a metagenomic dataset. The high-quality NCLDV and *Preplasmiviricota* genomes and core gene information recovered by BEREN can be used for multiple downstream applications. This includes finding interactions between these two groups of viruses and assessing their abundance and activity dynamics over time through metagenomic read mapping [53]. A dedicated tool to uncover the diversity of coexisting eukaryotic viruses in the *Bamfordvirae* kingdom in environmental sequence datasets will facilitate formulation of new hypothesis regarding virus-virus interactions and their roles in shaping the population of micro- and macroeukaryotes in diverse ecosystems.

## Data Availability

BEREN is an open-source tool available on Gitlab (https://gitlab.com/benminch1/BEREN/-/tree/master). All scripts to run the program, as well as test metagenomes are available there. Public metagenomes used for testing BEREN can be found using accessions in Table S1.

## Author contributions

BM and MM jointly developed the research idea. BM developed the pipeline and wrote most of the manuscript. MM supervised the research, secured funding and critically edited the manuscript.

## Acknowledgments

We gratefully acknowledge the use of the Pegasus supercomputer made available by the University of Miami Frost Institute of Data Science and Computing (IDSC).

