## Supplementary material for "BEREN: A bioinformatic tool for recovering Giant viruses, Polinton-like Viruses, and Virophages in metagenomic data": Figure S1

BEREN Supplemental

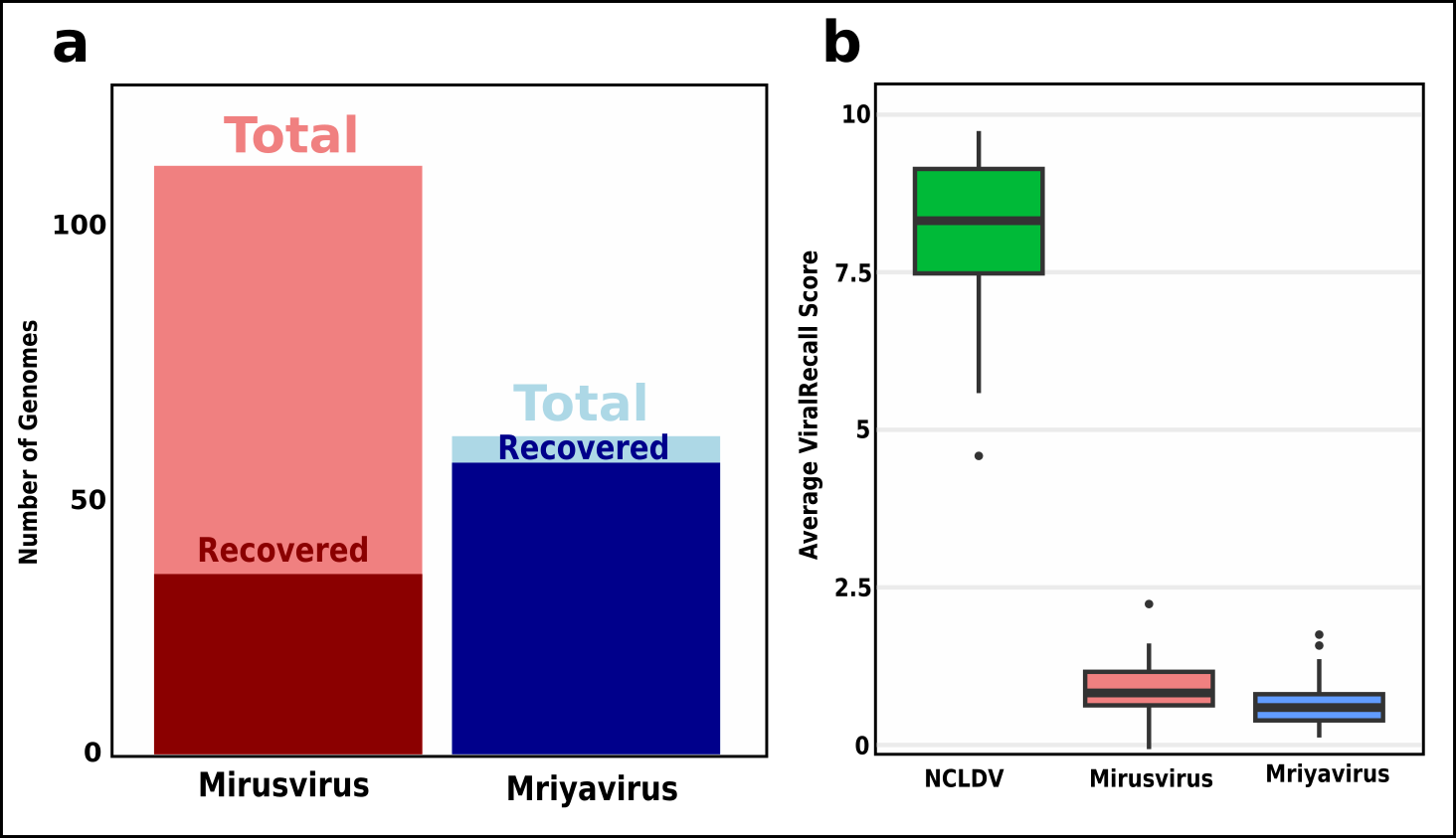

**Figure S1. BEREN’s ability to recover Mirusvirus and Mriyavirus genomes. (a)** A total of 111 Mirusvirus and 60 Mriyavirus genomes were used as input to BEREN as part of a mock metagenome. The total number of genomes recovered and identified by BEREN is shown above. **(b)** The average ViralRecall score for Mirusviruses and Mriyaviruses as compared to NCLDV genomes recovered using the BEREN pipeline.

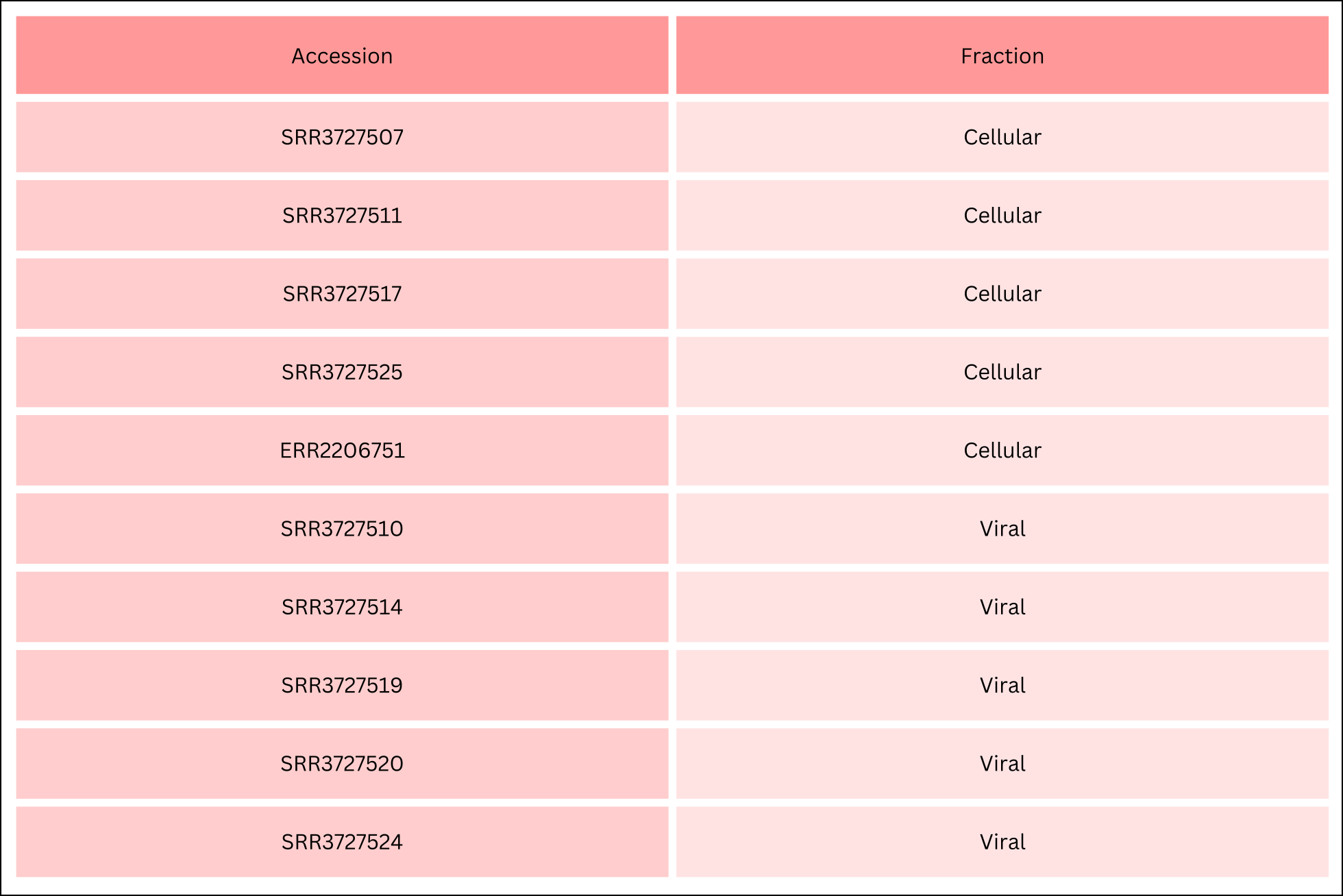

**Table S1. Baltic metagenome accessions for viral (<0.2uM) and cellular (>0.2uM) size fraction.**
